# Easternmost distribution of Asian hornet (*Vespa velutina nigrithorax*, du Buysson 1905) in Slovakia: Urgent need for advanced detection and interregional monitoring

**DOI:** 10.1101/2024.10.28.620586

**Authors:** Balázs Kolics, Éva Kolics, Zoltán Ács, Helena Mališová Proková, Katarína Baldaufová Senková, Dušan Senko

**Affiliations:** Festetics Bioinnovation Group, Georgikon Campus, Institute of Genetics and Biotechnology, Hungarian University of Agriculture and Life Sciences, H-8360, Keszthely, Hungary; Venic Nature Knowledge and Conservation Foundation, H-8400, Ajka, Hungary; Proková Medula s.r.o., Gercenova 6/b 85101, Bratislava, Slovakia; Department of Evolution and Systematics, Institute of Botany Plant Science and Biodiversity Centre, Slovak Academy of Sciences, Dúbravská cesta 9, SK-845 23 Bratislava, Slovakiaanický ústav SAV, Dúbravská cesta 9, 845 23 Bratislava, Slovakia

**Keywords:** first nest, Asian hornet, Slovakia, radio telemetry, molecular analysis, mtDNA, easternmost distribution

## Abstract

The invasive Asian hornet (*Vespa velutina nigrithorax*) continues its spread across Europe, posing a significant threat to biodiversity, viticulture, and apiculture. Species identification on the first individuals caught in Slovakia was confirmed via molecular analysis of the mitochondrial cytochrome c oxidase subunit I (COI) gene. Radio telemetry was employed to track foraging hornets, leading to the discovery of the nest within inaccessible private property. Mastering and utilizing manual tracking techniques assisted by radio telemetry, alongside public awareness campaigns and regional monitoring programs, are crucial in Slovakia and neighbouring countries to mitigate the potential ecological and economic impacts of this invasive species.

## INTRODUCTION

The invasive Asian hornet (*Vespa velutina nigrithorax* du Buysson, 1905), continues its relentless spread across Europe, posing a significant threat to biodiversity, viticulture and apiculture. Native to Southeast Asia, this social wasp predator has rapidly expanded its range since its accidental introduction to France in 2004 (Haxaire et al., 2006, Villemant et al., 2011, Villemant et al., 2006). Numerous European countries have reported established populations of *V. v. nigrithorax*, leading to concerns about its ecological and economic impact (Monceau et al., 2014, Arca et al., 2015). Slovakia, with its suitable climate and habitat conditions, has been identified as a potential area for *V. velutina* invasion (Fournier et al., 2017). The risk of establishment is further heightened by the ongoing expansion of this species in neighbouring countries, such as Hungary (Márta and Vas, 2023) and Czech republic (Walter et al., 2024) or Austria (Schorkopf et al., 2024). Early detection and rapid response are crucial for mitigating the potential damage caused by *V. velutina* (Roy et al., 2023).

## Materials and Methods Study Site

This study was conducted in Palárikovo, a village located in southwestern Slovakia. The first *Vespa velutina* specimen was collected on the 2^nd^ of October, 2024 in a private garden a banana plant (*Musa basjoo*) (Location 1). A second observation (48°01’57.3”N 18°04’21.6”E) occurred approximately 314 m away, where several individuals were observed foraging for insects on inflorescence of ivy (*Hedera helix*) (Location 2). These two locations served as the starting points for radio telemetry tracking.

### Nest location

To locate V. *velutina* nests, we employed radio telemetry using a Yupiteru MVT 7100 receiver (Yupiteru Industries Co., LTD.; Japan) and an iCOM R20 receiver (iCOM America Inc., USA), both equipped with YAGI-type directional antennas. Hornets were tagged following a modified version of the methodology described by Kennedy et al. (Kennedy et al., 2018), omitting the inactivation (chilling) step to ensure optimal flight capability. At Location 1, we used Dessau G19 D4 transmitters (weight ≤ 190 mg; operating frequency 150.042 MHz; operating time > 4 days; Plecotus Solutions GmbH, Germany). At Location 2, we used Dessau V5 transmitters (weight ≤ 250 mg; operating frequency 150.269 MHz; operating time > 10 days; Plecotus Solutions GmbH, Germany). Tracking was conducted on foot by following the signal direction and intensity until the hornet’s movement pattern suggested the likely location of a nest. Radio telemetry tracking was performed on October 2^nd^, 2024, under predominantly sunny conditions with a maximum daytime temperature of 17 °C.

### Molecular Genetic Analysis

To confirm species identification, we sequenced the mitochondrial cytochrome c oxidase subunit I (COI) gene. Three adult hornet specimens (females, workers) were collected in October 2024 in Palárikovo, Slovakia (48°02’04”N 18°04’14”E, 48°02’01”N 18°04’16”E). DNA was extracted from the posterior femoral muscles of each individual using the DNeasy Blood & Tissue Kit (Qiagen, Germany) following the manufacturer’s instructions. A 690 bp fragment of the COI gene was amplified via PCR using the universal barcoding primers HCO-2198 and LCO-1490 (Folmer et al., 1994). The thermocycling profile consisted of an initial denaturation step of 94°C for 3 minutes, followed by 36 cycles of 95°C for 1 minute, 40°C for 1 minute, and 72°C for 1 minute 30 seconds, with a final extension step of 72°C for 7 minutes.

PCR products were electrophoresed on a 1.5% TBE agarose gel, stained with ethidium bromide, and visualized under UV light. Amplicons were purified using the NucleoSpin Gel and PCR Clean-up Kit (Macherey-Nagel GmbH, Düren, Germany) according to the manufacturer’s protocol.

Sanger sequencing was performed in a 10 μl reaction volume containing 50-55 ng of purified PCR product and 10 pmol of each primer.

Sequences were analyzed and aligned using BioEdit software (Hall, 1999) and trimmed to 613-617 bp. To determine the phylogenetic relationship of the Slovakian hornet colony, its COI sequences were compared to *V. velutina nigrithorax* COI haplotypes deposited in the NCBI database (https://www.ncbi.nlm.nih.gov/). We included 31 *V. velutina* COI sequences representing all 18 known haplotypes (Takeuchi et al., 2017).

Phylogenetic analysis was conducted using the Maximum Likelihood method with the Tamura 3-parameter model (Tamura, 1992) in MEGA11 software (Tamura et al., 2021). The Tamura 3-parameter +G model was selected based on the lowest Bayesian Information Criterion (BIC) and Akaike Information Criterion (AIC) scores. Initial trees for the heuristic search were generated using Neighbor-Joining and BioNJ algorithms applied to a matrix of pairwise distances, with the topology exhibiting the superior log likelihood value selected. A discrete Gamma distribution was used to model evolutionary rate heterogeneity among sites. The final dataset comprised 34 nucleotide sequences with 613-617 positions.

## Results and Discussion

### Morphometrical and molecular confirmation

The easternmost record of *V. velutina* in Slovakia, discovered in Palárikovo on September 28^th^, 2024 (Purkart et al 2024), was confirmed by morphological identification of the captured specimens according to identification keys by Archer (Archer and Penney, 2012) was followed by molecular confirmation. All sequences obtained were deposited in GenBank under accession numbers PQ459388, PQ459391, and PQ459392. Comparison with NCBI reference sequences revealed 100% similarity with sequences from European populations (Fig. 1), confirming the species as *V. velutina nigrithorax*.

**Fig 1.**
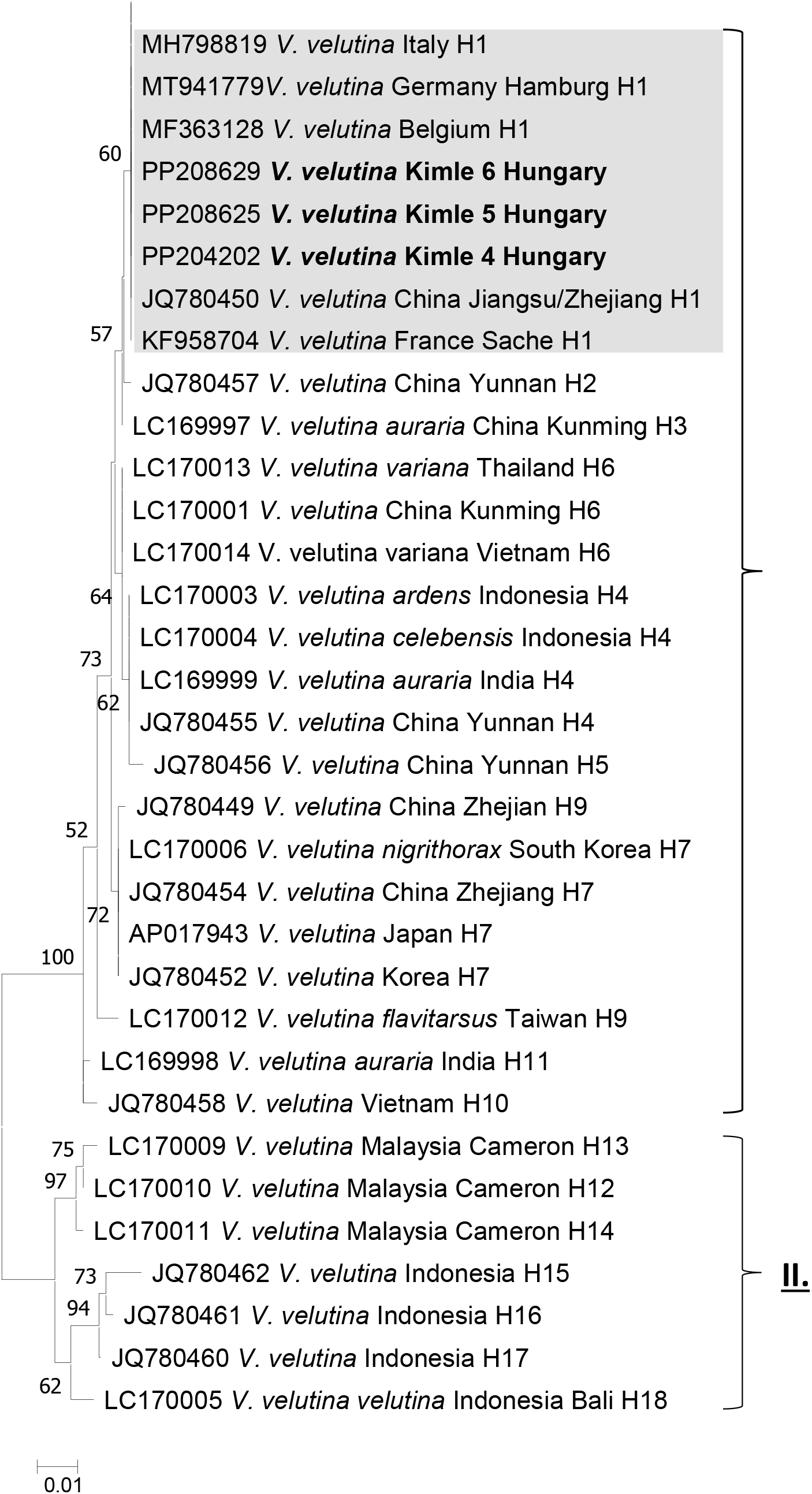
Maximum Likelihood phylogenetic tree constructed using the Tamura 3-parameter model. Branch support values are shown as percentages. The analysis involved 34 nucleotide sequences. Evolutionary analyses were conducted in MEGA11. The COI sequences of the three Slovakian hornet individuals share the same haplotype (H1) as other European populations and the sample from China. The COI haplotype numbers (H1–H18) follow (Takeuchi et al., 2017). *V. velutina* haplotypes divide into two distinct clades (I. and II.).

### V. v. nigrithorax Populations in Slovakia

The nest closest to Location 2 was located after unsuccessful triangulation, on the October 2^nd^, using tracking with two separate receivers equipped with directional antennas, triangulating its position using radio telemetry signal detection from two vantage points with an accuracy of 7 metres. Further tracking was precluded by the nest’s location within private property.

Having telemetry data, nest tracking was continued deploying thermal imaging drone to pinpoint its precise position on the following day. Favourable weather conditions (e.g., low wind speed, specific temperature range, 2.5 °C thermal difference) were critical to the success of this approach enabling that this thermal anomaly led to the visualisation of the first *V. velutina* nest in Slovakia (Purkart et al 2024). The inactivated nest exhibited a dark brownish coloration and the presence of several small holes, suggesting prior intrusion, possibly by birds. This observation, coupled with the fact that *V. velutina* typically exhibits strong nest defense and constructs its nests in exposed locations with a single entrance, suggests a higher probability of nest abandonment, which could have led to at least dispersal the queens.

According to repeated occurrence of two workers was observed foraging 1,359 and 1,640 m from the nest and a discovery of a male in Dolny Ohaj on October 15^th^, 2024, 14.5 km from the Palárikovo nest was observed (Purkart et al 2024).

The eradication of the Palárikovo nest in early October, similar to the removal of a nest in a likely isolated Hungarian population in 2023, may have helped to slow the spread of the invasive hornet. However, the relatively late discovery of the nest, along with finding in Dolny Ohaj could indicate that the species has established a broader presence in the region and is not confined to a single isolated location.

Previous detections in neighbouring countries, such as the August 2023 discovery of *V. velutina* workers in Kimle, Hungary (57.2 km from Palárikovo), in Pilsen in the Czech Republic V. *velutina* was reported on October 5^th^, 2023 and in August 7^th^, 2024 in the Northern part of Moravian region of the Czech Republic and the April 2024 observation of a founder queen in Salzburg, Austria, underscore the importance of vigilance and proactive monitoring across the region. This highlights the importance of continuous monitoring across wider geographical areas.

### Implications for Surveillance and Control of V. velutina in Central Europe

The presence of *V. velutina nigrithorax* population in the South of Slovakia marks a significant step in the eastward expansion of this invasive species. While the sporadic nature of its continental spread may still hold true in a broader context, the estimated speed of expansion (Bertolino et al., 2016, Verdasca et al., 2021, Robinet et al., 2017) calls for increased awareness in the Hungarian, Austrian and Slovak border area for the following year.

Although nest discoveries in Central Europe have been limited to date, the probability of further Asian hornet nests appearing in the region, will likely increase in 2025 as the species continues to spread.

This detection raises concerns about further undetected *V. velutina* populations in Slovakia and neighboring countries, highlighting the urgent need for regional monitoring programs, public awareness campaigns, and a clear legal framework for rapid response. This framework should streamline permissions for radio telemetry and drone-based nest detection, particularly concerning private property access and airspace regulations, to facilitate timely intervention.

This situation underscores the necessity of integrating advanced monitoring techniques, such as the combined radio telemetry approach employed in this study. Continued technological advancements in this field are essential, including enhancing the sensitivity and precision of radio telemetry for tracking the invasive hornet in diverse environments and refining the integration of radio telemetry with drone technology. These improvements will enable more efficient and targeted deployment of resources, facilitating the timely detection and eradication of *V. velutina*.

## Acknowledgements

We would like to thank to the Hungarian Carnica Association and Kolics Apiaries for providing transmitters and the receivers for the experiment.

## Ethical Statement

None.

## Funding Statment

This research work was supported by the Hungarian Government and the European Union, with the co-funding of the European Regional Development Fund in the frame of Széchenyi 2020 Programme GINOP2.3.2-15-2016-00054.

## Data availability

the data on which the study is based were generated during the course of this study. Raw data are available from the corresponding author upon request.

## Authors’ Contributions

H.M.P.: Radio telemetry tracking, data analysis, manuscript review, and compilation of information regarding *V. velutina*; Z.Á.: Phylogenetic analysis and manuscript preparation.

É.K.: Manuscript preparation, molecular investigations, radio telemetry tracking; B.K.: Manuscript preparation, molecular investigations, radio telemetry tracking, and overall coordination of the research. D.S.: Initial discovery of *Vespa velutina*, member of the scientific expert committee for eradication efforts in Palárikovo, and manuscript preparation; K.B.S.: Initial discovery of *Vespa velutina*

## Specimen Deposition

Voucher specimens will be deposited in the following institutions:

- Hungarian University of Agriculture and Life Sciences, Keszthely Campus (in ethanol)

